# Automatic Detection and Neurotransmitter Prediction of Synapses in Electron Microscopy

**DOI:** 10.1101/2021.11.02.467022

**Authors:** Angela Zhang, S. Shailja, Cezar Borba, Yishen Miao, Michael Goebel, Raphael Ruschel, Kerrianne Ryan, William Smith, B.S. Manjunath

## Abstract

This paper presents a deep-learning based workflow to detect synapses and predict their neurotransmitter type in the primitive chordate *Ciona intestinalis* (*Ciona*) EM images. Identifying synapses from electron microscopy (EM) images to build a full map of connections between neurons is a labor-intensive process and requires significant domain expertise. Automation of synapse detection and classification would hasten the generation and analysis of connectomes. Furthermore, inferences concerning neuron type and function from synapse features are in many cases difficult to make. Finding the connection between synapse structure and function is an important step in fully understanding a connectome. Activation maps derived from the convolutional neural network provide insights on important features of synapses based on cell type and function. The main contribution of this work is in the differentiation of synapses by neurotransmitter type through the structural information in their EM images. This enables prediction of neurotransmitter types for neurons in *Ciona* which were previously unknown. The prediction model with code is available on Github.

## Introduction

We propose a deep learning convolutional neural network to detect *Ciona* intestinalis synapses from serial section transmission electron microscopy (EM) images, as well as a multi-modal method of predicting neurotransmitter type based on several modalities of data obtained in different ways - EM imaging, light microscopy of *in situ* hybridization, and behavioral observation experiments. *Ciona* intestinalis is of interest to neuroscientists because of its close evolutionary relationship to vertebrates and small nervous system. Moreover, it is one of the very few animals for which a complete synaptic connectome is available [1]. An essential aspect of constructing a connectome, a comprehensive map of neurons and their connections in the brain, is to identify the synapses which form chemical and electrical communication links between neurons [17]. Traditionally, detection of synapses is completed manually through expert analysis of thousands of electron microscopy (EM) images. This is a time consuming process that can take thousands of hours for one specimen. For example, the first *Ciona* connectome was constructed manually over a period of 5 years from serial section EM images [1]. Another crucial but missing component of understanding the connectome is the classification of the function and properties of each synapse. One of the most important properties is neurotransmitter use.

While some experimental methods such as *in situ* hybridization can identify cells, or clusters of cells, expressing transcripts indicating neurotransmitter use [2], the resolving power of light microscopy cannot reveal the connectivity of these neurons. Additionally, in cases with intermingled neurotransmitter types and high variability in neuron location across specimens, it may not be possible to find correspondence between neuron properties derived by situ hybridization and neurons identified by serial EM in the course of constructing the connectome. Visual inspection of the synapses in *Ciona* EM images do not reveal any distinguishable differences between known excitatory and inhibitory neurons. Some examples are included in Figure 1. Like all chemical synapses, those in *Ciona* include the vesicle cluster in the presynaptic neuron, which varies in count and size, and the postsynaptic density, which also varies in size and density. No apparent pattern in these features is visible upon manual inspection.

**Figure 1.**
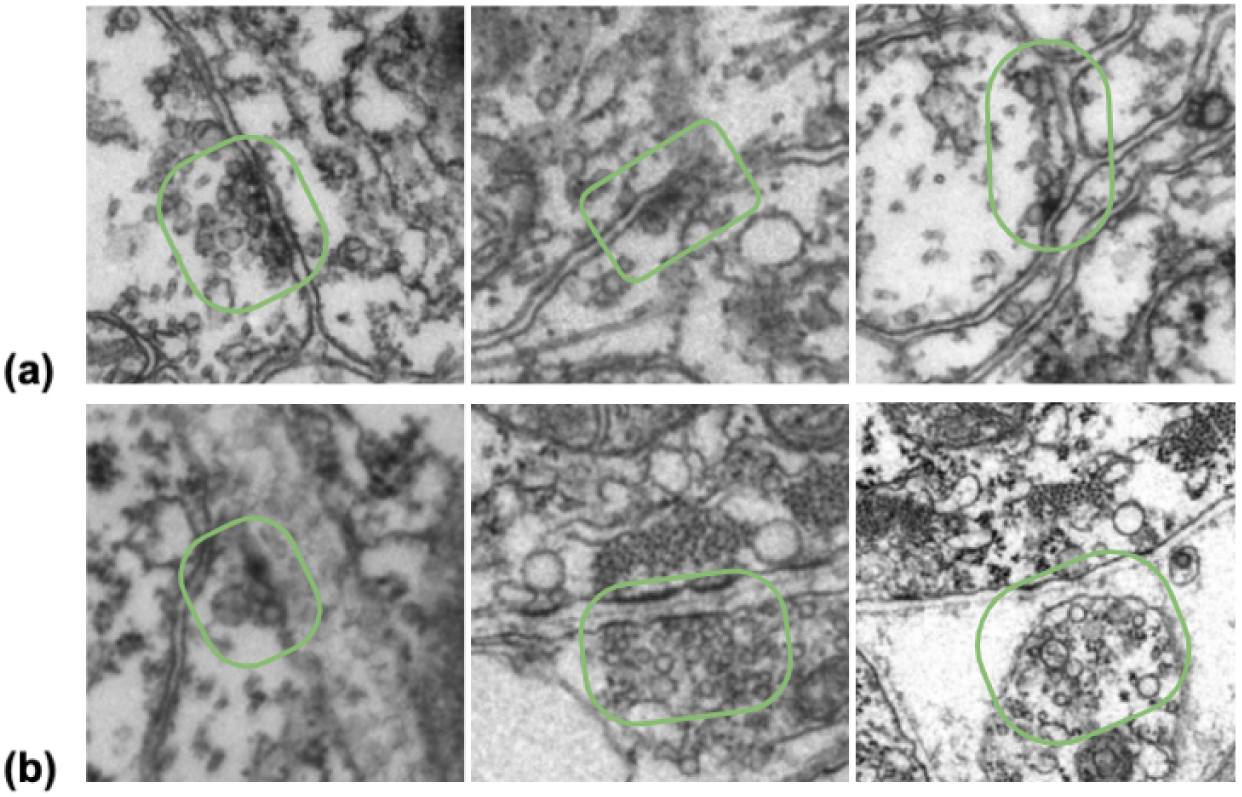
Examples of (a) inhibitory and (b) excitatory synapses. The synaptic region is circled in green. It can be seen that there are varying vesicle counts and sizes, as well as little visible post-synaptic density.

**Figure 2.**
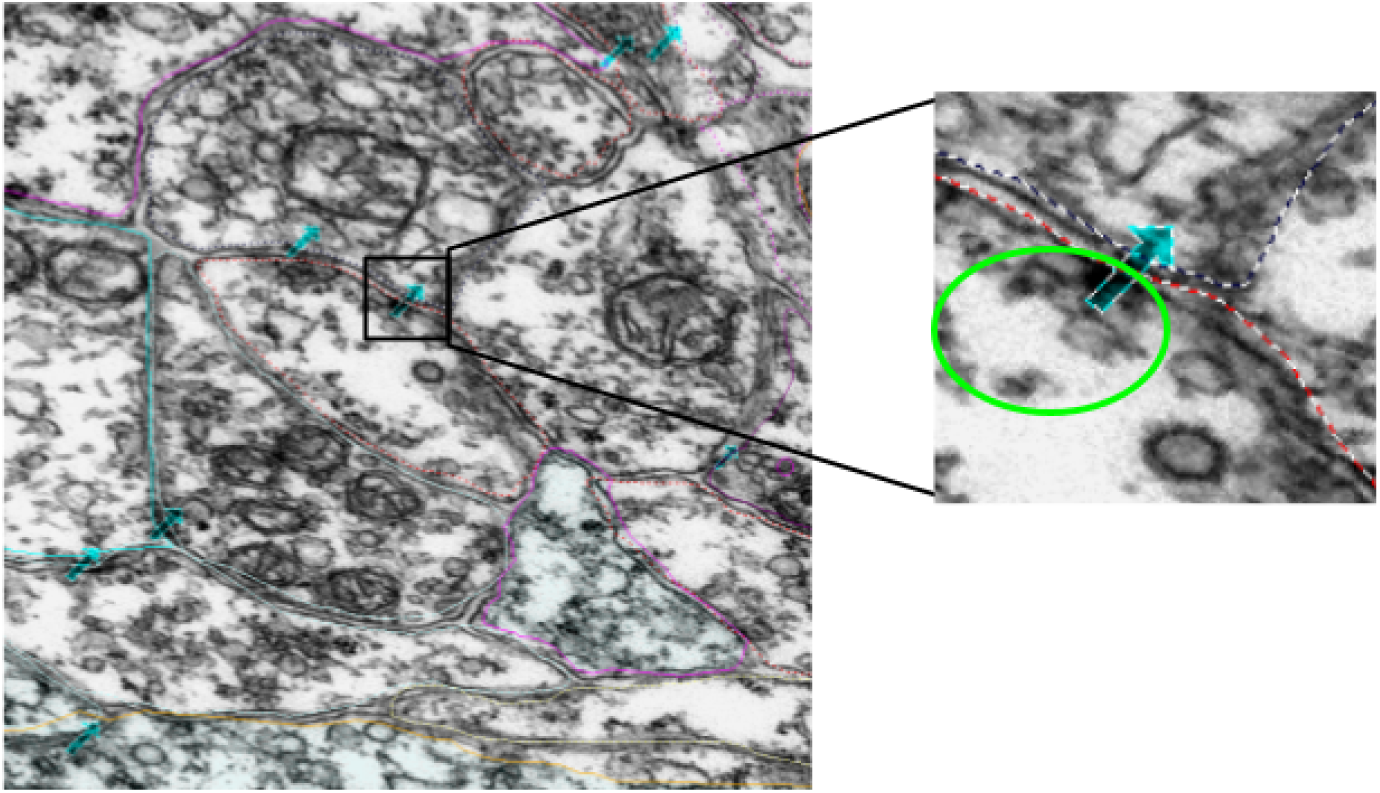
Example of manual annotations of synapses, indicated by the cyan arrows. The zoomed in image shows a patch containing a synapse, with the vesicles encircled with bright green and the cell boundaries marked with dotted lines. The direction of the arrow does not indicate synaptic direction.

In this paper, we use manually annotated EM images of a *Ciona* larva to validate our findings. We also propose a deep learning method to predict the neurotransmitter type of a neuron based on the appearance of its synapses in EM images, and generate activation maps from the prediction model to visualize the features which are important in synapse classification.

### Contributions

Our main contributions are as follows.

1. An automated method to detect and localize synapses from EM data with high accuracy.
2. A method to predict the neurotransmitter class of neurons in *Ciona* based on its synapse structure. Creation and analysis of class activation maps (CAM) from the neural network to derive the synaptic features which are identified as important to neurotransmitter type prediction.
3. 3 Model-based predictions of neurotransmitter class by cell which are previously unknown, and can be used in further experiments to help determine the true neurotransmitter expression of said cells.

The methods and data are available upon request.

## Prior Work

Previous work have reported successful application of computer vision methods for automatically detecting synapses in EM images of *Drosophila*, mouse and rabbit neurons [3, 4, 5]. However, the synapses of these organisms contain unique features which the detection systems rely on heavily. The algorithm for *Drosophila* synapse detection [3] uses primarily the t-shaped feature and postsynaptic density to detect the synapse with a 3D UNet [6], while the algorithms for mouse synapse detection [4], [5] was created for cryogenic EM. One approach applies pixel-level classification and graph cut segmentation on EM images to identify potential synapses, then filters the potential synapses with a random forest classifier [4]. The classifiers are trained on manually annotated samples. A second approach finds handcrafted synaptic cleft features for the presynaptic and postsynaptic regions, and uses LogitBoost to perform the final synapse detection [5]. A third publication describes the uses of a fusion of ribbon, cleft, and vesicle features of a rabbit retina synapse to detect retinal synapses through kernel learning [7]. Ribbons are not ubiquitous amongst synapses and clefts are not always visible for different types of synapses, so this method would not work on all types of synapses.

Studies have qualitatively shown differences between excitatory and inhibitory synapses - namely that the vesicle shape and post-synaptic density appearance varied between the two functions [8, 9, 10]. However, these papers are mostly qualitative, and do not provide predictive functionality. Furthermore, these studies do not analyze a large number of synapses, and are not directly translatable to *Ciona* synapses due to differences between species. More recently, synapse classification using a deep network was done on EM images of *Drosophila* neurons [11]. This body of work is most similar to the one presented in this chapter, but the appearance of Drosophila synapses differs drastically from those of *Ciona*. Our previous study [2] combined in situ hybridization with the existing connectome derived from EM to determine the neurotransmitter expression of neuron types, such as the photoreceptors. Point cloud matching was done to match relative cell locations in 3 dimensions between fluorescent microscopy and electron microscopy results. While we are successful in determining the neurotransmitter expression of individual cells belonging to the photoreceptors, we are unable to determine with certainty the neurotransmitter expression of individual cells belonging to the relay neurons and other types. The present study aims to take steps towards resolving the neurotransmitter assignments in these ambiguous regions while applying neural network approaches to this problem.

## Data

The data consisted of EM images from a total of 3375 60 nm serial sections from the anterior brain vesicle to the motor ganglion of a *Ciona* larva. The sections were collected and imaged at 3.85 nm per pixel [1]. The data collected surpasses 1 terabyte. The original *Ciona* EM serial section image data set was collected, processed and annotated using the program RECONSTRUCT [12]. The annotations we focused on are the synapse annotations, which are stored as points in 3 dimensions with a naming system to indicate the pre- and post-synaptic neurons. In order to facilitate analysis, python scripts were written to interface with the program and stored data and extract image patches corresponding to annotated regions. A series of geometric transforms was used to co-register the annotation coordinates with the aligned 3D image stack coordinates. The RECONSTRUCT images used in this project are available upon request. The image processing and extraction steps are described in the methods section.

### Data Curation and Annotation

The EM images were scaled and aligned in the z-dimension while annotating the cellular components. Image scaling is specified by a pixel size (magnification) parameter while alignment is represented by a non-linear transformation associated with the image. Each transformation maps trace points or image pixels into the section using a combination of basis functions representing an elementary motion such as translation, orientation, scaling, and deformation. We extract the underlying annotations (actual section coordinates in microns) from each section by combining these movement components in different proportions. Each image is associated with an independent transformation which determines the size and location of the element on the section. Applying the inverse transform on the contour point (x,y), we obtain the points (x’, y’), on which applying the forward image transformation brings the points to the original image domain. For our study, synapse is represented by seven points forming an arrow as shown in Figure 3. After getting the coordinates of these points on the original image domain, we determine the centroid of these points, and extract a 500×500 dimensional patch around this centroid point. We perform this operation for all the synapse annotations to obtain approximately 25,000 total number of patches. The code to extract synapse points from EM data can be found at https://github.com/s-shailja/Ciona_EM.

**Figure 3.**
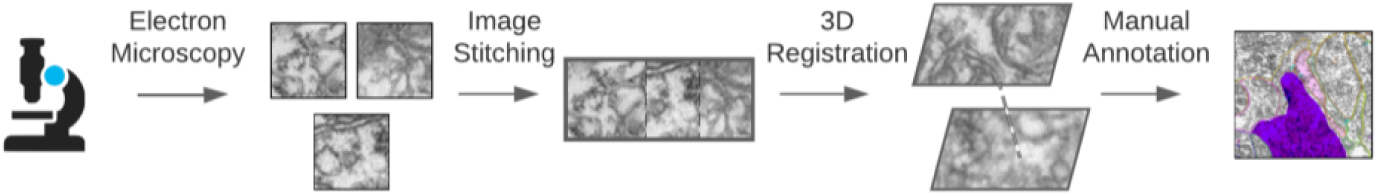
Illustration of Data Preparation Method.

## Methods

The major steps involved in *Ciona* synapse detection and classification are shown in Figure 4. The synapse detection workflow includes image patch extraction and training of a ResNeXT network. The neurotransmitter prediction workflow is similar, but sorts the images based on the assigned IDs [1] of the presynaptic cell before training the deep learning network. Post-prediction class activation maps of the convolutional neural network are computed for better understanding of important imaging features for classification. Feature maps are reduced to 2 dimensional space and plotted to visualize the feature-space distance between synapses of various neurotransmitter types.

**Figure 4.**
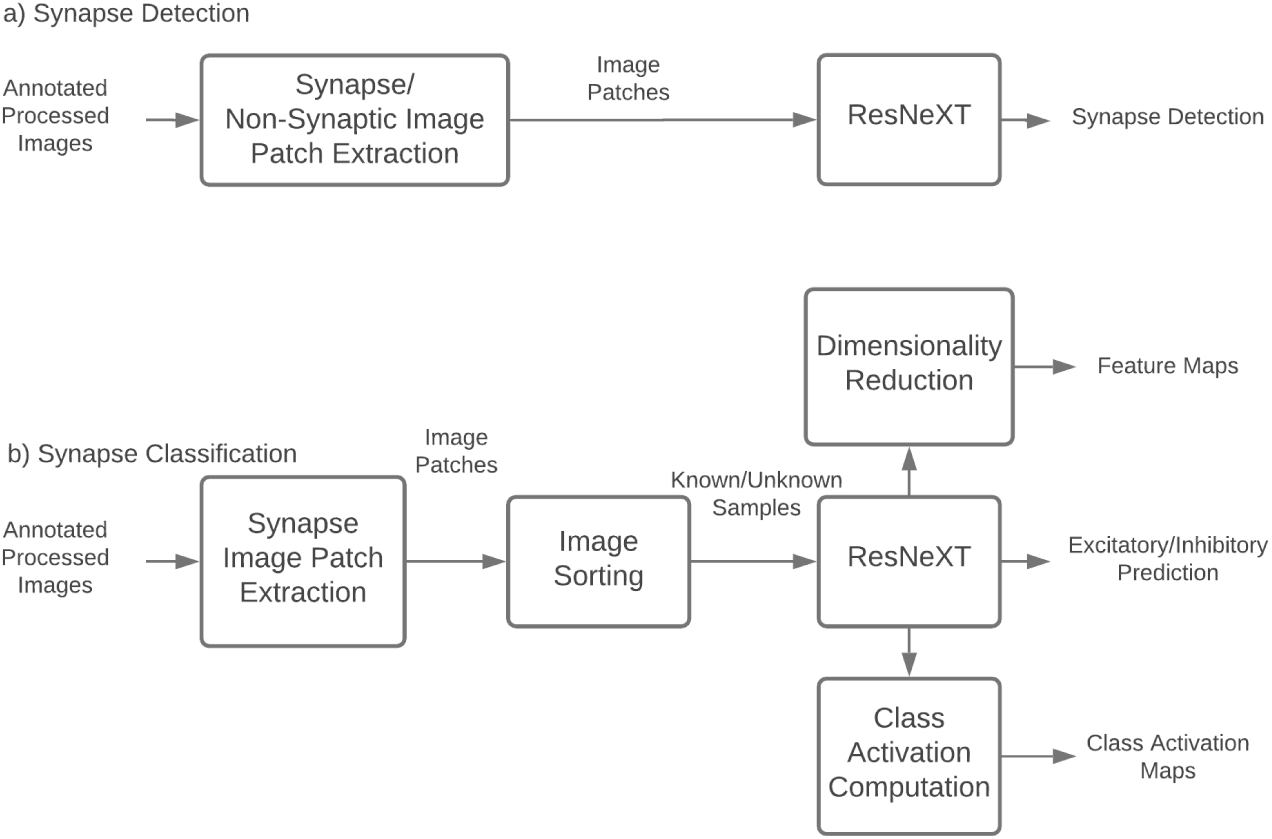
Synapse Detection / Classification flowchart.

For synapse detection and classification, the same training process is used, as follows. A ResNeXt-50 network architecture [13] pretrained on the ImageNet [14] dataset was retrained on the extracted image patches. First, the last fully connected layer was replaced with a randomly initialized fully connected layer with an input of 2048 and an output of 2. The output has 2 possible classes, with 0 being inhibitory and 1 being excitatory. The training was done in two sessions. First, all layers of the ResNeXt are “frozen”, with no gradient, except for the last fully connected layer. The model is trained for 100 epochs, and the best model is used for the second round of training. On the second round of training, the entire network is “unfrozen”, every layer of the network is tuned with retraining for 200 epochs. The architecture of the deep network is shown in Table 1.

**Table 1.** ResNeXt-50 architecture.

| stage | layers |  | output |
| --- | --- | --- | --- |
| conv1 | 7x7, 64 | stride 2 | 112x112 |
| conv2 (x3) | 3x3 max pool | stride 2 | 56x56 |
|  | 1x1, 128 | C = 32 |  |
|  | 3x3, 128 |  |  |
|  | 1x1, 256 |  |  |
| conv3 (x4) | 1x1, 256 | C = 32 | 28x28 |
|  | 3x3, 256 |  |  |
|  | 1x1, 512 |  |  |
| conv4 (x6) | 1x1, 512 | C = 32 | 14x14 |
|  | 3x3, 512 |  |  |
|  | 1x1, 1024 |  |  |
| conv5 (x3) | 1x1, 1024 | C = 32 | 7x7 |
|  | 3x3, 1024 |  |  |
|  | 1x1, 2048 |  |  |
|  | global average pool |  | prediction (0 or 1) |
|  | 1000-d fc, softmax |  |  |

### Synapse Detection

Image patches containing annotated synapses and image patches containing non-synaptic structures were used for the training (80%) and testing (20%) sets. 1,400 500×500 image patches containing synapses, and 1,392 500×500 image patches containing non-synaptic structures, were used for training a synapse detection network. Non-synaptic structures used were annotated variously as botrysomes, coated vesicles, basal bodies, and autophagosomes. Annotated structures which were avoided due to the possibility of synapses being in the same patch included terminals, vesicles, dense core vesicles, and gap junctions. Misclassified patches are analyzed visually for confounding factors.

### Synapse Neurotransmitter Prediction

Image patches were grouped by presynaptic and postsynaptic cell ID [1] before splitting into training, validation, and testing sets. A similarity computation was applied to each patch to ensure that no duplicates or extremely overlapping patches were used. Due to the limited number of synapses and known neuron groups available, we combined the two inhibitory neurotransmitter types (glycine, gamma-Aminobutyric acid (GABA)) and the two excitatory neurotransmitter types (glutamate, acetylcholine). 470 excitatory synapses and 338 inhibitory synapses were used for training a synapse detection network. 1246 excitatory synapses and 396 inhibitory synapses were used for testing. Each synapse was composed of 3-10 image patches in the z dimension.

Nine neuron groups with 4 known neurotransmitter types, as determined by *in situ* hybridization [2], were used in training and testing. Each presynaptic neuron had 1-41 associated synapses. Neurotransmitter class was predicted using the trained network on an additional 13 neuron groups with previously unknown neurotransmitter types. For each presynaptic neuron, a majority vote was made from the predictions of each synapse belonging to the neuron. Based on the strength of consensus, network confidence, and prior knowledge from *in situ* hybridization experiments [2], predictions were made on the neurotransmitter type of neurons which surpassed a confidence interval described as follows. For each neuron, we tallied the number of predictions for each neurotransmitter valence. This tally is referred to as a vote. The valence with the most votes is chosen as the raw prediction of that presynaptic neuron. If the votes for each valence are close (e.g., inhibitory vote is not more than *e*^1.02^ times of the excitatory votes) or that the total number of votes is less than 3, we determine the prediction to be inconclusive.

### Class Activation Maps

512×512 image patches with labels are passed through the neurotransmitter prediction neural network. The final layer of our network uses global average pooling to go from a size of 16×16×2048 to 2048 features, then compute probabilities for each of the N classes using a fully connected layer without bias. For each pixel in the 16×16 feature map, we then compute the amount that this feature contributes to the output class. We apply the fully-connected weights to the features at each pixel, cutting out the average pooling step. The basic idea for deriving class activation maps is described in [15]. After extracting the activation maps from the model, we use connected component analysis to find seed points, then employ a watershed algorithm to segment the activation map into disjoint regions to further analyze spatial and intensity information about the activations. Next, we removed regions that were smaller than a set threshold, which was set empirically to 5000 pixels. Following this step, for each remaining connected region, we then calculated the x and y coordinates of the centroid for the region and the average activation intensity inside the region of interest.

### Feature Maps

To better understand the network’s decision boundaries, the high-dimensional features from the output layer of the prediction network were reduced to 20 dimensions using Principal Component Analysis. The 20 dimensional feature is then further reduced to 2 dimensions using t-distributed Stochastic Neighbor Embedding [16], which better visualizes the clustering characteristics of the features. The features are grouped in various ways to gain more insight into how they are clustered.

## Results

For synapse detection, a training accuracy of 0.99 and a testing accuracy of 0.98 was achieved. For training, 2780 samples were used, with half of the samples containing synapses and half without synapses. For testing, 692 samples were used, also with a 50/50 split of synapse vs. non-synapse sample. There were no false negatives, but there were 13 false positives which were detected by the model. Examples of some false positives are shown in Figure 5.

**Figure 5.**
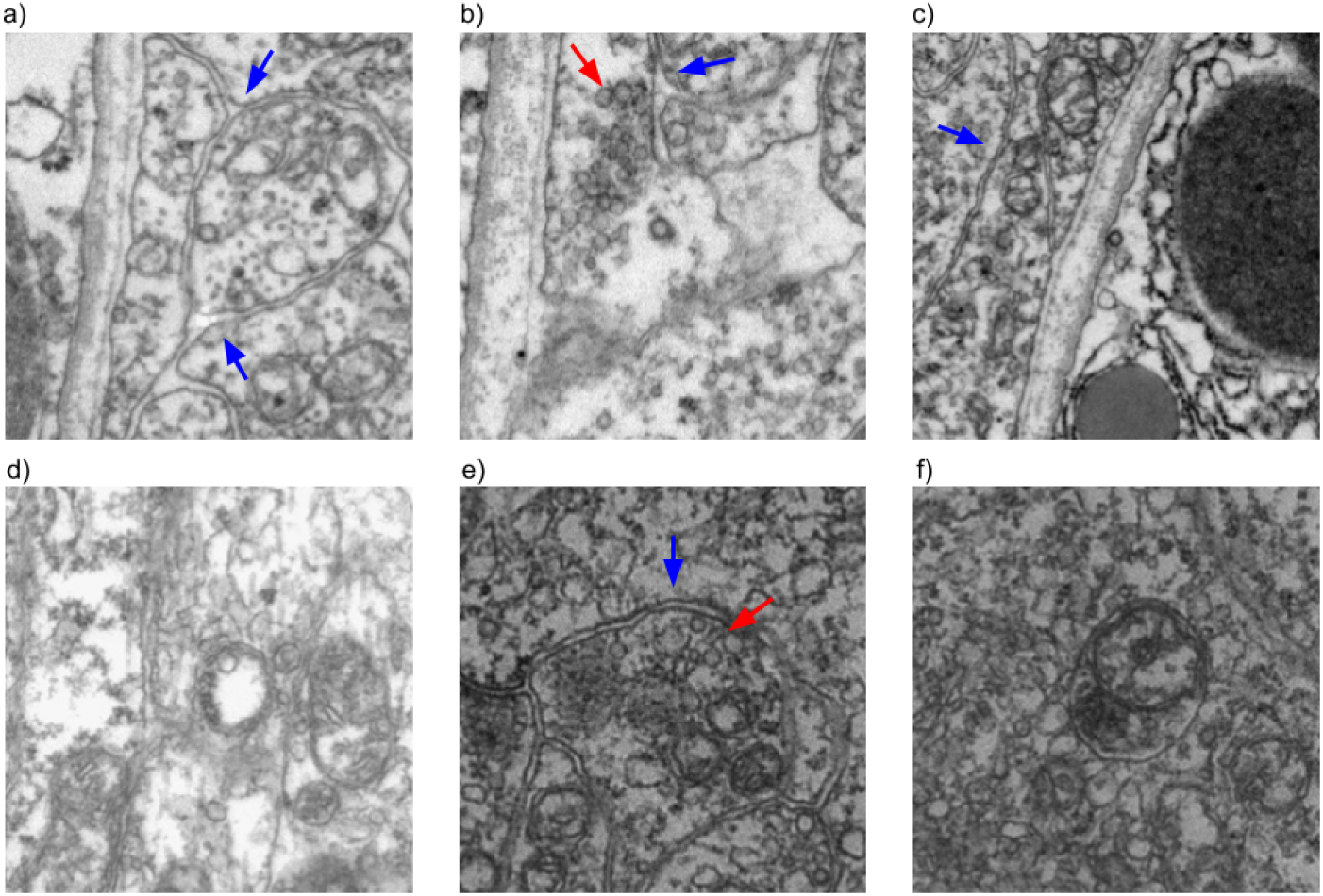
Examples of false positives for the synapse detection network. a), b), and c) contain coated vesicles, d) and e) contain botrysome, and f) contains an autophagosome. It can be seen that either cell boundaries (blue arrows) or groups of vesicles (red arrows) are visible in many of the cases.

Performance of the network on classification is shown in T and Table 2. Precision is the number of True Positives (TP) divided by the sum of TP and True Negatives (TN), and recall is TP divided by the total number of positive samples. Precision determines how often selected items are relevant, and recall determines how often relevant items are selected.

**Table 2.**
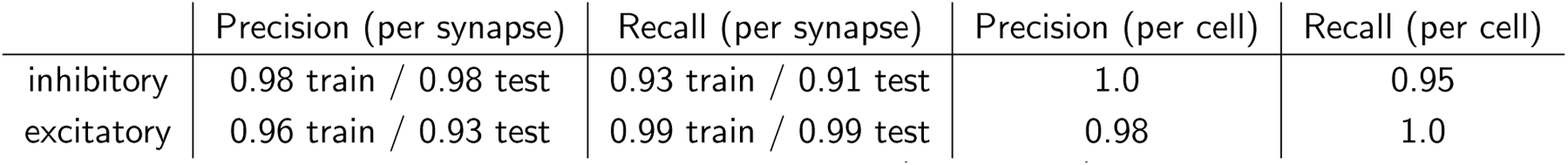
Automated synapse classification performance - per synapse (train and test) and per cell.

|  | Precision (per synapse) | Recall (per synapse) | Precision (per cell) | Recall (per cell) |
| --- | --- | --- | --- | --- |
| inhibitory | 0.98 train / 0.98 test | 0.93 train / 0.91 test | 1.0 | 0.95 |
| excitatory | 0.96 train / 0.93 test | 0.99 train / 0.99 test | 0.98 | 1.0 |

**Table 3.** Automated synapse classification performance breakdown by neurotransmitter type. As seen in the table, the performance for glycine was the worst, most likely due to the low number of cells and synapses available to train the model. Performance for GABA, acetylcholine, and glutamate were similar to each other.

|  | Number of Cells | Number of Synapses | Accuracy (per cell) | Accuracy (per synapse) |
| --- | --- | --- | --- | --- |
| gly | 3 | 22 | 0.67 | 0.71 train / 0.6 test |
| gaba | 19 | 457 | 1.0 | 0.92 train / 0.95 test |
| ach | 19 | 292 | 1.0 | 0.97 train / 1.0 test |
| glut | 23 | 342 | 1.0 | 1.0 train / 0.98 test |
| overall | 64 | 1113 | 0.98 | 0.95 train / 0.97 test |

Of the failed detections (all false positives), 10 cases were image patches annotated as coated vesicles, 2 cases were annotated as botrysomes, and 1 case annotated as an autophagosome. Some representative patches are included in Figure 5. From the failed cases, it can be seen that they tend to contain cell boundaries and vesicles, which are features associated with synapses. The lack of false negatives is reassuring, as the goal of the detection network is to detect likely synapses for screening by experts, so false positives are better tolerated than false negatives.

To get a better idea of the features which are important to the neurotransmitter predictor network, we visualized the activation maps of the network on different classes, as described in the Methods section. The results of the visualization show that cell boundaries, vesicles, and postsynaptic density are the main focus of the majority of the attention for the trained network. Some examples of activation maps are shown in Figure 6

**Figure 6.**
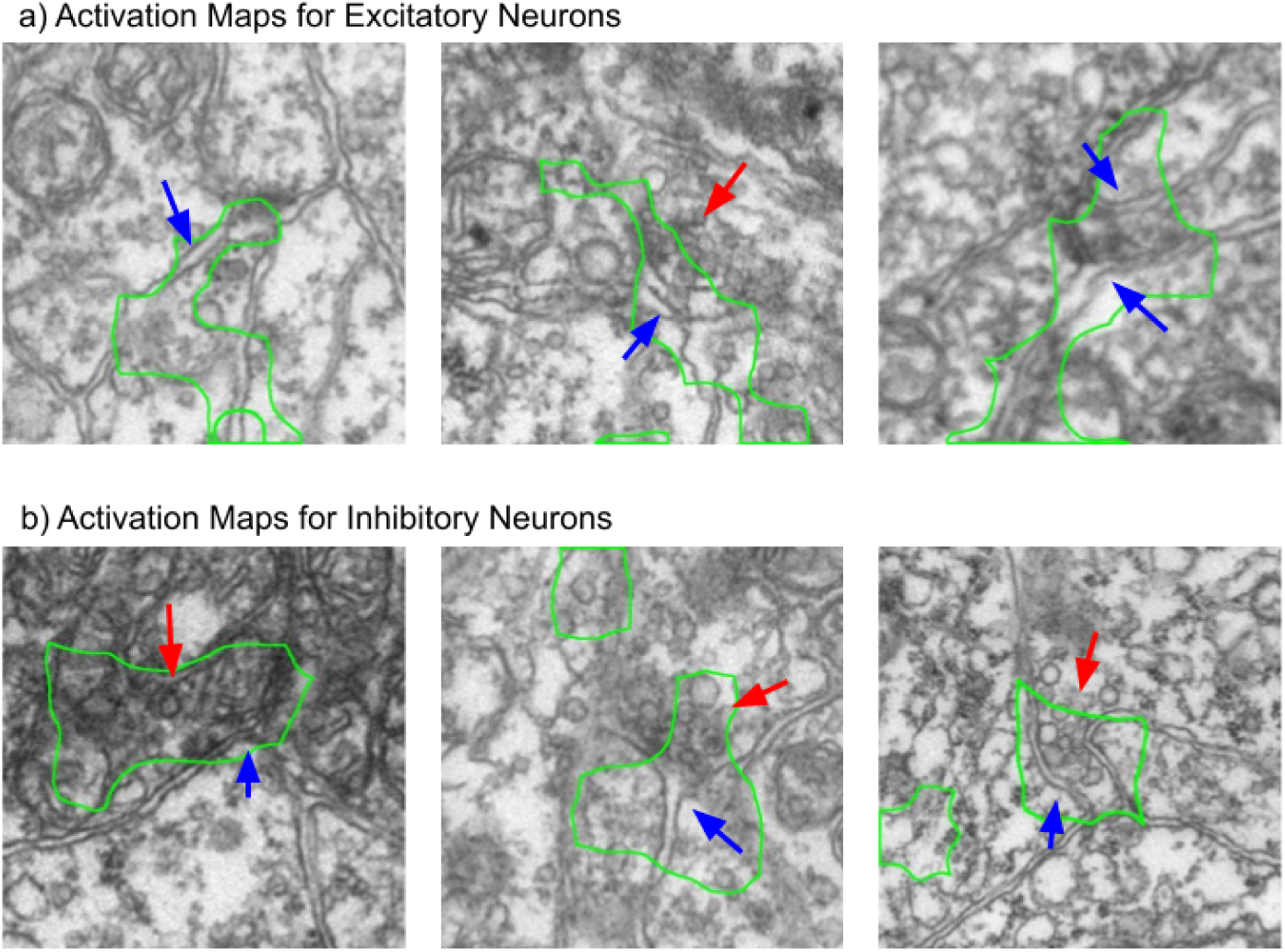
Processed Class Activation Maps for Excitatory and Inhibitory Synapse Image Patches. The network seems to pay more attention to the vesicles when predicting inhibitory neurons, and the cell boundary when predicting excitatory neurons. The blue arrows indicate cell boundaries while the red arrows indicate vesicles. The green outline shows the main region of interest of the network.

The feature maps derived from the model outputs are shown in Figures 7 - 9. From the feature maps, it is evident that the synapses are placed in clusters which mostly correspond to their neurotransmitter class, inhibitory or excitatory. From Figure 9 it appears that the differences between different excitatory neurotransmitters (acetylcholine and glutamate) are also captured by the model, even though this information was not explicitly included during training. From Figure 7, it can be seen that the features of synapses tend to be spread out throughout the feature space, and co-mingle amongst cell groups. This is promising, because this indicates that the model is picking up on differences between synapses which are more indicative of neurotransmitter class, rather than group-specific differences.

**Figure 7.**
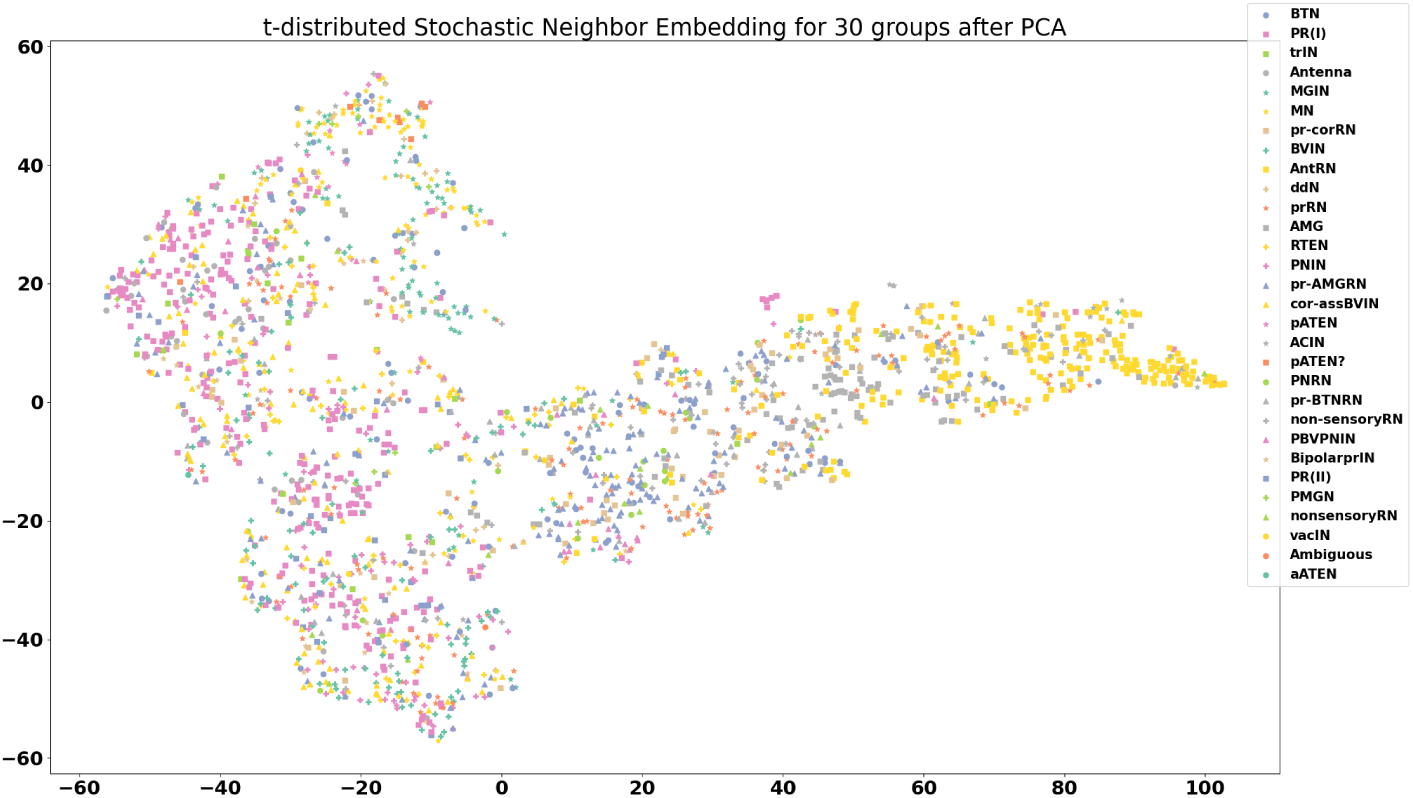
Visualization of features for 30 neuron groups after reduction to 2 dimensions using principal component analysis and t-distributed stochastic neighbor embedding. Each data point on the plot is the computed feature of a synapse, with synapses that span multiple frames averaged across all frames. There is quite a bit of intermingling of the features between cell groups, which is an encouraging sign that the model is picking up on differences not unique to each cell type.

**Figure 8.**
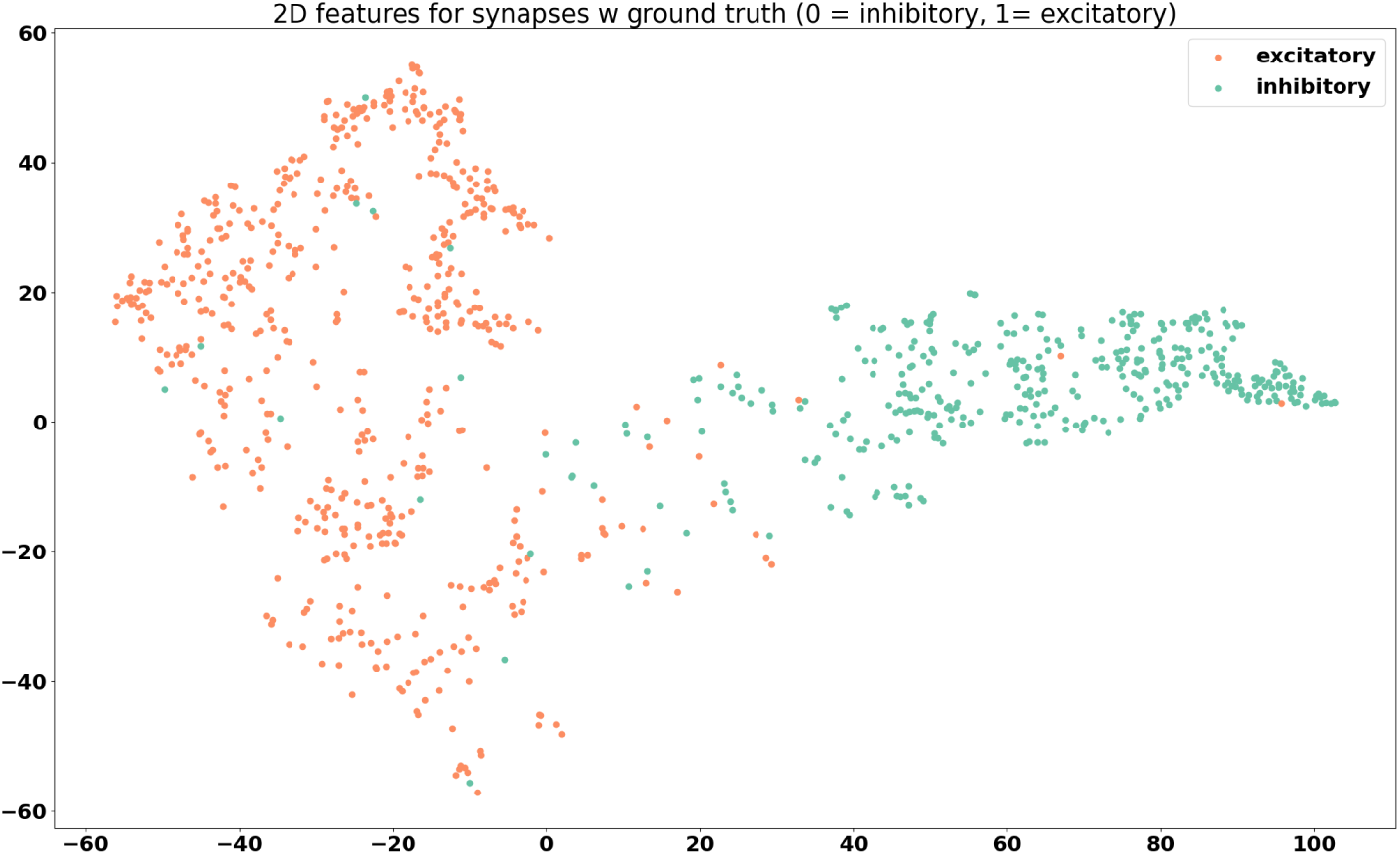
Feature visualization for synapses belonging to cells with known neurotransmitter type, grouped by valence. Two distinct groups can be seen.

**Figure 9.**
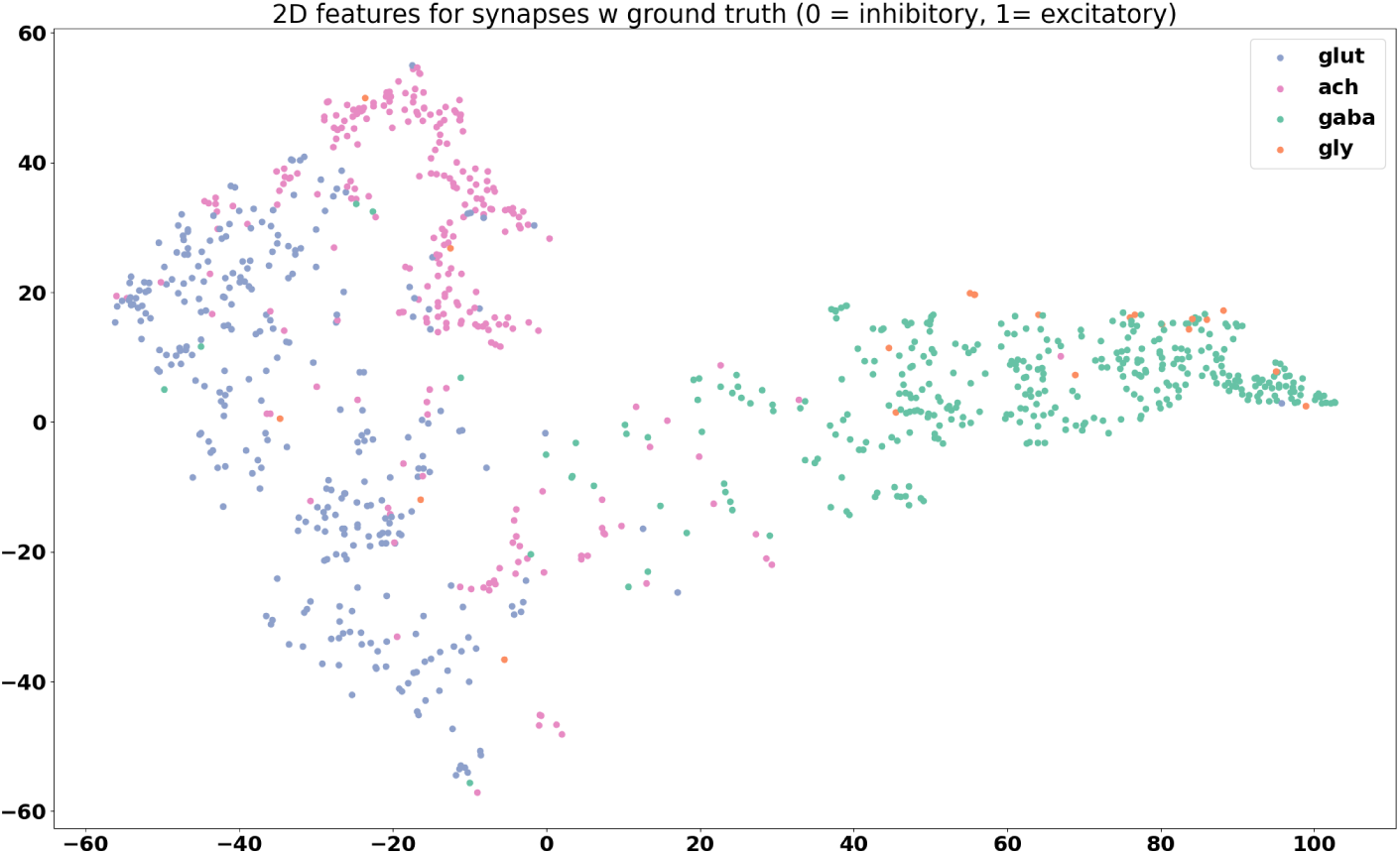
Feature visualization for synapses with known neurotransmitter type, grouped by neurotransmitter. Even though the prediction model was trained only to differentiate between excitatory and inhibitory synapses, it seems that the features tend towards separation by neurotransmitter type, with the exception of glycine, again likely do to the lack of training samples.

Table 5 shows the performance of the model on synapses belonging to cells of known neurotransmitter type. Some of these synapses were used for training, and others were used for testing. We tallied the number of predictions by presynaptic neuron type and valence (inhibitory and excitatory) in the training set. The tally is repeated for the testing set. The tally for each valence is referred to as a vote. The valence with the most votes is chosen as the raw prediction of that presynaptic neuron. Those are the values in the Predicted Valence section. We compared the valence of the raw prediction with the neurotransmitter type determined by *in situ* hybridization to calculate precision and recall.

Table 4 shows the predictions of the model on relay neurons of unknown neurotransmitter type. We noticed that the prediction model tends towards predicting more excitatory synapses in the relay neuron group, since the observed average number of excitatory relay neurons from *in situ* hybridization was 11, while the number of predicted excitatory relay neurons was 14 [2]. All but one of the pr-AMG relay neurons was predicted to be excitatory, with varying degrees of likelihood. This is different from our predictions in [10], which indicated an inclination towards inhibitory pr-AMG relay neurons, but with low confidence. However, the predicted number of excitatory and inhibitory neurons in the relay neuron group is closer to the observed number in [2] than the predicted results from 3D point cloud matching, which suggests a promising direction for resolving the neurotransmitter types in that region. Further experiments and analysis is needed to determine with certainty the neurotransmitter type of each relay neuron.

**Table 4.**
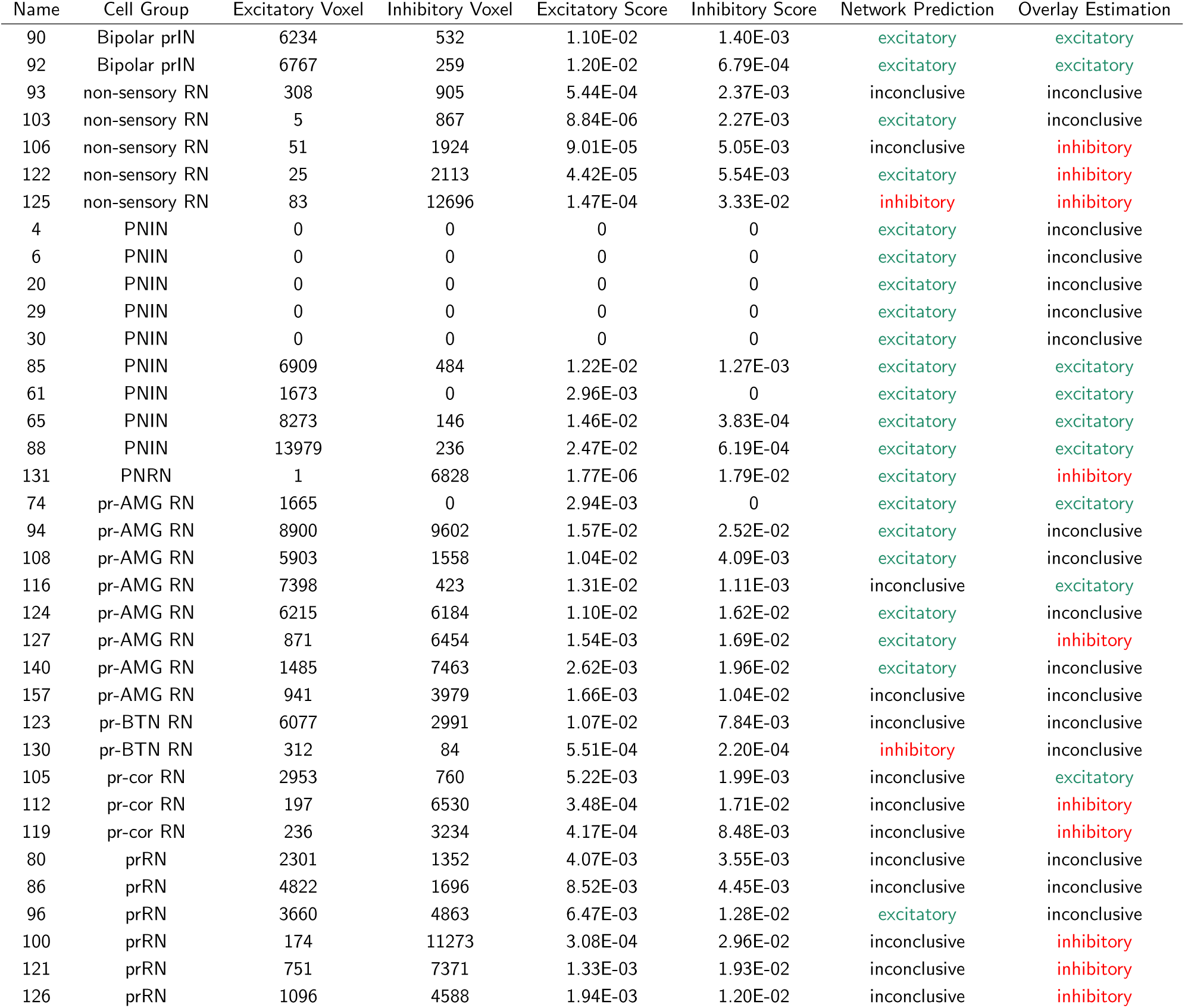
Predicted neurotransmitter valence for the relay neurons (RNs). RNs used for training with known neurotransmitter type are omitted from this table and included in the appendix. The column with our model predictions is ‘Network Prediction’, which is ‘inconclusive’ if there are fewer than 3 synapses for a given cell, or if the number of excitatory and inhibitory predictions are similar, that is, one is not more than 1.5 times the other.

| Name | Cell Group | Excitatory Voxel | Inhibitory Voxel | Excitatory Score | Inhibitory Score | Network Prediction | Overlay Estimation |
| --- | --- | --- | --- | --- | --- | --- | --- |
| 90 | Bipolar prIN | 6234 | 532 | 1.10E-02 | 1.40E-03 | excitatory | excitatory |
| 92 | Bipolar prIN | 6767 | 259 | 1.20E-02 | 6.79E-04 | excitatory | excitatory |
| 93 | non-sensory RN | 308 | 905 | 5.44E-04 | 2.37E-03 | inconclusive | inconclusive |
| 103 | non-sensory RN | 5 | 867 | 8.84E-06 | 2.27E-03 | excitatory | inconclusive |
| 106 | non-sensory RN | 51 | 1924 | 9.01E-05 | 5.05E-03 | inconclusive | inhibitory |
| 122 | non-sensory RN | 25 | 2113 | 4.42E-05 | 5.54E-03 | excitatory | inhibitory |
| 125 | non-sensory RN | 83 | 12696 | 1.47E-04 | 3.33E-02 | inhibitory | inhibitory |
| 4 | PNIN | 0 | 0 | 0 | 0 | excitatory | inconclusive |
| 6 | PNIN | 0 | 0 | 0 | 0 | excitatory | inconclusive |
| 20 | PNIN | 0 | 0 | 0 | 0 | excitatory | inconclusive |
| 29 | PNIN | 0 | 0 | 0 | 0 | excitatory | inconclusive |
| 30 | PNIN | 0 | 0 | 0 | 0 | excitatory | inconclusive |
| 85 | PNIN | 6909 | 484 | 1.22E-02 | 1.27E-03 | excitatory | excitatory |
| 61 | PNIN | 1673 | 0 | 2.96E-03 | 0 | excitatory | excitatory |
| 65 | PNIN | 8273 | 146 | 1.46E-02 | 3.83E-04 | excitatory | excitatory |
| 88 | PNIN | 13979 | 236 | 2.47E-02 | 6.19E-04 | excitatory | excitatory |
| 131 | PNRN | 1 | 6828 | 1.77E-06 | 1.79E-02 | excitatory | inhibitory |
| 74 | pr-AMG RN | 1665 | 0 | 2.94E-03 | 0 | excitatory | excitatory |
| 94 | pr-AMG RN | 8900 | 9602 | 1.57E-02 | 2.52E-02 | excitatory | inconclusive |
| 108 | pr-AMG RN | 5903 | 1558 | 1.04E-02 | 4.09E-03 | excitatory | inconclusive |
| 116 | pr-AMG RN | 7398 | 423 | 1.31E-02 | 1.11E-03 | inconclusive | excitatory |
| 124 | pr-AMG RN | 6215 | 6184 | 1.10E-02 | 1.62E-02 | excitatory | inconclusive |
| 127 | pr-AMG RN | 871 | 6454 | 1.54E-03 | 1.69E-02 | excitatory | inhibitory |
| 140 | pr-AMG RN | 1485 | 7463 | 2.62E-03 | 1.96E-02 | excitatory | inconclusive |
| 157 | pr-AMG RN | 941 | 3979 | 1.66E-03 | 1.04E-02 | inconclusive | inconclusive |
| 123 | pr-BTN RN | 6077 | 2991 | 1.07E-02 | 7.84E-03 | inconclusive | inconclusive |
| 130 | pr-BTN RN | 312 | 84 | 5.51E-04 | 2.20E-04 | inhibitory | inconclusive |
| 105 | pr-cor RN | 2953 | 760 | 5.22E-03 | 1.99E-03 | inconclusive | excitatory |
| 112 | pr-cor RN | 197 | 6530 | 3.48E-04 | 1.71E-02 | inconclusive | inhibitory |
| 119 | pr-cor RN | 236 | 3234 | 4.17E-04 | 8.48E-03 | inconclusive | inhibitory |
| 80 | prRN | 2301 | 1352 | 4.07E-03 | 3.55E-03 | inconclusive | inconclusive |
| 86 | prRN | 4822 | 1696 | 8.52E-03 | 4.45E-03 | inconclusive | inconclusive |
| 96 | prRN | 3660 | 4863 | 6.47E-03 | 1.28E-02 | excitatory | inconclusive |
| 100 | prRN | 174 | 11273 | 3.08E-04 | 2.96E-02 | inconclusive | inhibitory |
| 121 | prRN | 751 | 7371 | 1.33E-03 | 1.93E-02 | inconclusive | inhibitory |
| 126 | prRN | 1096 | 4588 | 1.94E-03 | 1.20E-02 | inconclusive | inhibitory |

The full prediction table is included in the Appendix, in Table 5. Summaries of the available data are shown in Figures 10 and 11.

**Figure 10.**
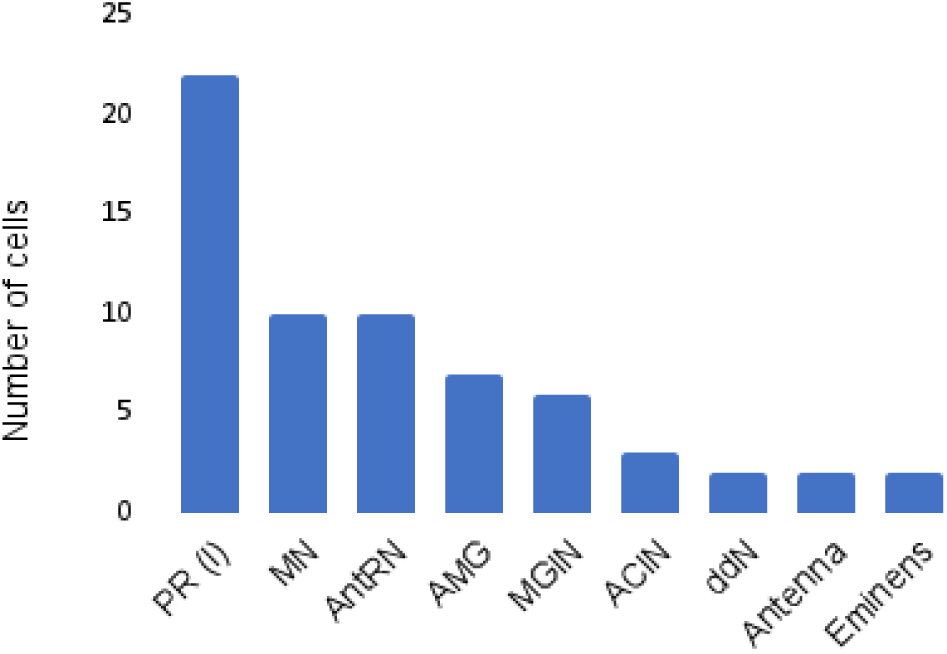
Number of unique cells per cell type of selected groups of interest.

**Figure 11.**
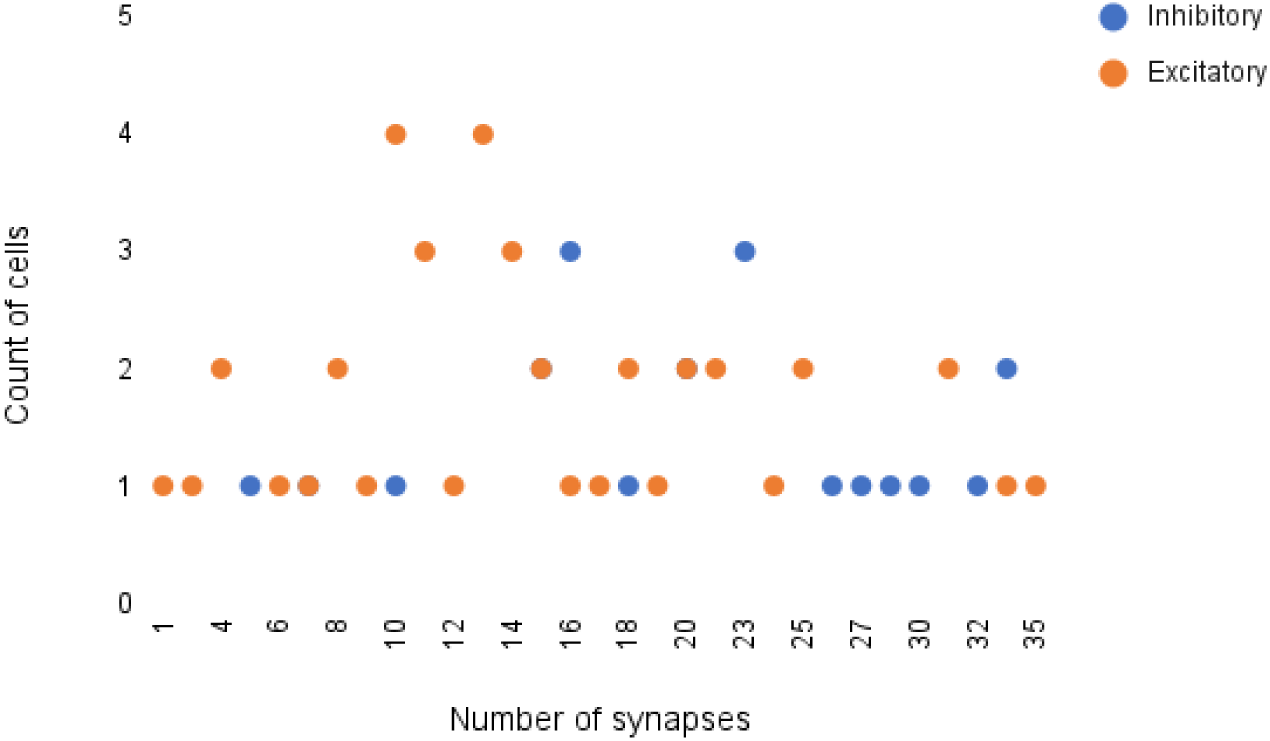
Frequency of cells with various number of synapses. It can be seen that the majority of cells have between 10-20 synapses.

### Comparison with Manual Overlay

The crux of our analysis lies in our hypothesis that the same cells appear in similar locations across *Ciona* specimens. To provide an additional point of comparison, we manually overlay the fluorescent imaging results from *in situ* hybridization with the cell centroids provided by the annotated EM data, as seen in Figure 12. Using the Unity software [17], the centroid and volume of each neuron in the four selected cell groups as given in [1] is rendered in 3D. Each neuron is approximated with a sphere of its corresponding volume. As seen on the leftmost image in Figure 9, image stacks of *in situ* from [2] which contain expression of both vesicular GABA transporter (VGAT, inhibitory) and vesicular Acetylcholine transporter (VAChT, excitatory) in the posterior Brain Vesicle are also rendered into the software with real-world dimensions. Since certain parts of the VGAT structure are well known and consistent, such as in the photoreceptors and at the posterior end of the relay neurons, this was used to manually align the *in situ* with the connectome. The matching criteria is as follows: the posterior border doesn’t extend beyond the most posterior antRN, the dorsal cap marks the Eminen cells, and the two patches, a smaller posterior one and larger anterior one, on the right mark the two photoreceptor groups, PR-I (only pr9 and pr10) and PR-II, respectively. After this alignment is done, the smaller VACHT-labelled regions are brought into view for analysis. Seven *in situs* from [2] are aligned using the mentioned structures and similarity across the *in situs* as guides. Once they are aligned, a collision detector is used to compute the number of voxels in contact with each neuron. An illustration of the matching process is shown in Figure 12, and the comparative results are shown in Table 4.

**Figure 12.**
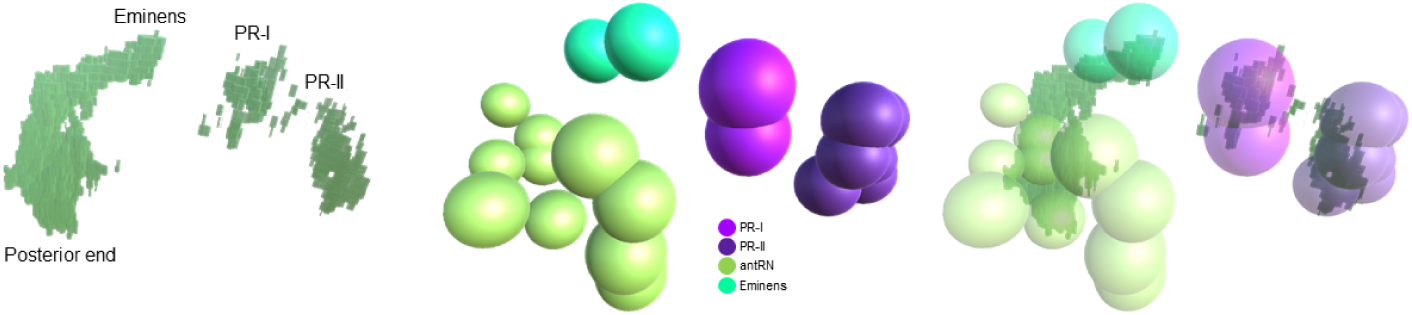
Illustration of manual overlay of in-situ hybridization results for VGAT (left image) and EM-derived cell centroids (middle image) in 3D. Of the four cell groups shown, The areas with VGAT are encapsulated by the cell models, as seen in the rightmost image. See text for more details.

## Discussion

The proposed synapse detection model has good performance on the detection of synapses in electron microscopy for *Ciona*. While there are some false positives from the model prediction (Table 1), this is desirable compared to false negatives, because an expert can then screen the predictions to determine true synapses. The ratio of synapses to non-synaptic structures in a typical EM of *Ciona* is on the order of 1:1000. Instead of scanning an entire image for possible synapses, the model can drastically reduce the annotation time needed for synapse annotation. The synapse prediction model has helped to identify possible neurotransmitter types for cells from certain neuronal groups, which were previously unknown. While we cannot be absolutely certain that the model has predicted correctly for synapses belonging to cells with previously unknown neurotransmitter types, the output of the model seems reasonable given the feature analysis we have done. Comparison with previous *in situ* hybridization results have also shown that the prediction of the model is likely correct. For the relay neurons, in [2] we had previously found an average of 16 VGAT-positive neurons and 11 VACHT-positive neurons. The prediction of the network matches these numbers well, and much better than the point-cloud registration method used in [2]. We hope that the model predictions will work in conjunction with both prior and future analysis to help resolve the neurotransmitter type of individual neurons in regions which have been difficult to resolve using in-situ hybridization and other experimental methods.

More work can be done on the analysis of cellular and subcellular features in neurons with undetermined neurotransmitter type. Overall connectivity and cell shape can be obtained from the annotated EM data and may be useful tools in better understanding the relationship between structural features and neurotransmitter type expression. The connection between cell group and synapse structure can also be further explored, and may help with a more robust neurotransmitter prediction method.

## Appendix

**Table 5.** Ground Truth Predictions.

| cell id | *in situ* | llr test | llr train | excitatory test | inhibitory test | excitatory train | inhibitory train | prediction test | prediction train |
| --- | --- | --- | --- | --- | --- | --- | --- | --- | --- |
| ACIN1L | gly | 0.24 | -1.73 | 1 | 0 | 0 | 6 | excitatory | inhibitory |
| ACIN2L | gly | -0.68 | 0.12 | 1 | 2 | 4 | 3 | inhibitory | excitatory |
| ACIN2R | gly | -0.46 | -0.54 | 0 | 1 | 1 | 3 | inhibitory | inhibitory |
| AMG1 | gaba | -1.08 | -1.66 | 0 | 5 | 1 | 9 | inhibitory | inhibitory |
| AMG2 | gaba | -0.67 | -3.09 | 1 | 4 | 1 | 14 | inhibitory | inhibitory |
| AMG3 | gaba | -0.21 | -3.59 | 0 | 1 | 0 | 14 | inhibitory | inhibitory |
| AMG4 | gaba | -0.47 | -2.42 | 0 | 2 | 2 | 12 | inhibitory | inhibitory |
| AMG5 | ach | 2.49 | 2.87 | 8 | 0 | 14 | 2 | excitatory | excitatory |
| AMG6 | gaba | -0.49 | -3.55 | 1 | 2 | 1 | 14 | inhibitory | inhibitory |
| AMG7 | gaba | -0.6 | -3.1 | 0 | 3 | 1 | 16 | inhibitory | inhibitory |
| Ant1 | glut | 0.21 | 3.89 | 3 | 1 | 14 | 0 | excitatory | excitatory |
| Ant2 | glut | 2.46 | 7.25 | 9 | 0 | 25 | 1 | excitatory | excitatory |
| 120 | gaba | -1.56 | -11.75 | 0 | 4 | 1 | 29 | inhibitory | inhibitory |
| 134 | gaba | -1.58 | -6.74 | 0 | 5 | 1 | 20 | inhibitory | inhibitory |
| 135 | gaba | -1.15 | -5.12 | 0 | 3 | 1 | 19 | inhibitory | inhibitory |
| 142 | gaba | -1.54 | -7.14 | 0 | 5 | 2 | 23 | inhibitory | inhibitory |
| 143 | gaba | -1.75 | -6.55 | 0 | 5 | 3 | 20 | inhibitory | inhibitory |
| 147 | gaba | -2.94 | -7.12 | 0 | 7 | 0 | 20 | inhibitory | inhibitory |
| 152 | gaba | -2.58 | -5.79 | 0 | 8 | 3 | 21 | inhibitory | inhibitory |
| 153 | gaba | -1.52 | -3.36 | 0 | 5 | 1 | 10 | inhibitory | inhibitory |
| 159 | gaba | -1.79 | -5.6 | 0 | 5 | 2 | 16 | inhibitory | inhibitory |
| 161 | gaba | -3.02 | -5.22 | 0 | 6 | 1 | 16 | inhibitory | inhibitory |
| ddNL | ach | 1.39 | 4.69 | 4 | 0 | 16 | 0 | excitatory | excitatory |
| ddNR | ach | 1.37 | 3.79 | 4 | 0 | 12 | 0 | excitatory | excitatory |
| Em1 | gaba | -1.11 | -7.92 | 1 | 6 | 3 | 31 | inhibitory | inhibitory |
| Em2 | gaba | -1.6 | -6.26 | 1 | 6 | 3 | 24 | inhibitory | inhibitory |
| MGIN1L | ach | 0.87 | 6.27 | 5 | 0 | 25 | 1 | excitatory | excitatory |
| MGIN1R | ach | 0.43 | 6 | 3 | 0 | 22 | 0 | excitatory | excitatory |
| MGIN2L | ach | 0.82 | 4.85 | 3 | 0 | 14 | 0 | excitatory | excitatory |
| MGIN2R | ach | 2.01 | 3.88 | 6 | 0 | 14 | 1 | excitatory | excitatory |
| MGIN3L | ach | 0.33 | 2.91 | 1 | 0 | 7 | 0 | excitatory | excitatory |
| MGIN3R | ach | 0.68 | 2 | 2 | 0 | 7 | 1 | excitatory | excitatory |
| MN1L | ach | 1.13 | 5.77 | 3 | 0 | 18 | 0 | excitatory | excitatory |
| MN1R | ach | 2.9 | 3.85 | 8 | 0 | 16 | 1 | excitatory | excitatory |
| MN2L | ach | 1.84 | 3.46 | 4 | 0 | 9 | 0 | excitatory | excitatory |
| MN2R | ach | 1.56 | 4.81 | 4 | 0 | 11 | 0 | excitatory | excitatory |
| MN3L | ach | 0.43 | 1.91 | 1 | 0 | 5 | 0 | excitatory | excitatory |
| MN3R | ach | 0.85 | 2.77 | 3 | 0 | 7 | 0 | excitatory | excitatory |
| MN4L | ach | 1.91 | 1.02 | 6 | 0 | 4 | 0 | excitatory | excitatory |
| MN4R | ach | 0.76 | 2.92 | 2 | 0 | 7 | 0 | excitatory | excitatory |
| MN5L | ach | 0.4 | 1.34 | 1 | 0 | 3 | 0 | excitatory | excitatory |
| MN5R | ach | 0.37 | 2.54 | 1 | 0 | 6 | 0 | excitatory | excitatory |
| pr1 | glut | 1.5 | 2.05 | 4 | 0 | 9 | 0 | excitatory | excitatory |
| pr11 | glut | 0.67 | 2.41 | 3 | 0 | 10 | 0 | excitatory | excitatory |
| pr12 | glut | 0.85 | 4.72 | 3 | 0 | 16 | 0 | excitatory | excitatory |
| pr13 | glut | 1.1 | 2.96 | 3 | 0 | 11 | 0 | excitatory | excitatory |
| pr14 | glut | 0.16 | - | 1 | 0 | 0 | 0 | excitatory | excitatory |
| pr15 | glut | 1.98 | 7.33 | 7 | 0 | 27 | 0 | excitatory | excitatory |
| pr16 | glut | 0.89 | 1.83 | 3 | 0 | 7 | 0 | excitatory | excitatory |
| pr17 | glut | 0.44 | 3.35 | 2 | 0 | 13 | 0 | excitatory | excitatory |
| pr18 | glut | 0.14 | 0.58 | 1 | 0 | 2 | 0 | excitatory | excitatory |
| pr19 | glut | 1.38 | 1.48 | 5 | 0 | 7 | 0 | excitatory | excitatory |
| pr2 | glut | 0.39 | 1.3 | 2 | 0 | 6 | 0 | excitatory | excitatory |
| pr20 | glut | 2.92 | 5.94 | 9 | 0 | 22 | 0 | excitatory | excitatory |
| pr21 | glut | 0.43 | 2.18 | 2 | 0 | 9 | 0 | excitatory | excitatory |
| pr22 | glut | 0.58 | 2.4 | 3 | 0 | 10 | 0 | excitatory | excitatory |
| pr23 | glut | 1.06 | 4.16 | 4 | 0 | 14 | 0 | excitatory | excitatory |
| pr3 | glut | 0.54 | 3.06 | 2 | 0 | 12 | 0 | excitatory | excitatory |
| pr4 | glut | 0.66 | - | 3 | 1 | 0 | 0 | excitatory | excitatory |
| pr5 | glut | 0.85 | 2.41 | 3 | 0 | 8 | 0 | excitatory | excitatory |
| pr6 | glut | 0.75 | 2.14 | 2 | 0 | 9 | 0 | excitatory | excitatory |
| pr7 | glut | 1.18 | 4.82 | 4 | 0 | 16 | 0 | excitatory | excitatory |
| pr8 | glut | 0.66 | 2.7 | 3 | 0 | 11 | 0 | excitatory | excitatory |
| pr9 | gaba | -0.36 | -1.3 | 0 | 1 | 4 | 11 | inhibitory | inhibitory |

## Acknowledgements

Thanks to Dr. Ian Meinertzhagen for his continued support of the project.

## Availability of data and materials

Data and materials are available upon request.

## Consent for publication

All authors have provided consent for publication.

## Authors’ contributions

Angela Zhang conceptualized the idea, developed the models and algorithms, and wrote the paper. S. Shailja extracted the data from RECONSTRUCT. Cezar Borba developed the method to compare expected results using the manual overlay. Yishen Miao developed the uncertainty measurements and created the tables. Michael Goebel and Raphael Ruschel developed the code for Class Activation Maps. Kerrianne Ryan conducted the experiments and created the annotations which provided the data for this paper. William Smith and B.S. Manjunath supervised the project and provided directional advice.

